# Myofibers drive postnatal myonuclear accretion through Arp2/3-dependent membrane protrusions

**DOI:** 10.64898/2026.03.22.713469

**Authors:** Catarina Sequeira, Silvia Di Francescantonio, Gabriele Motta, Naoko Kogata, Michael Way, Edgar R. Gomes

**Author notes:** corresponding author/lead contact: Edgar R. Gomes Gulbenkian Institute for Molecular Medicine Faculdade de Medicina da Universidade de Lisboa Av. Professor Egas Moniz 1649-028 Lisboa Portugal. These authors contributed equally to this work (Catarina Sequeira, Silvia Di Francescantonio).

## Abstract

Skeletal muscle postnatal growth depends on myoblast fusion with pre-existing myofibers, but whether myofibers actively regulate this process remains unclear. The Arp2/3 complex, which nucleates branched actin networks, is required for myoblast fusion during embryogenesis, but its role in postnatal myofibers is unknown. Here, we investigated the role of the Arp2/3 complex in myofibers using an inducible skeletal muscle–specific Arpc4 knockout model. We find that loss of Arp2/3 in myofibers impairs myonuclear accretion, resulting in smaller myofibers and reduced muscle strength. Satellite cells are still activated and able to differentiate but accumulate around Arpc4-deficient myofibers rather than fusing, pointing to a myofiber-intrinsic defect. In co-culture, myofibers form long-lived membrane protrusions enriched in Arp2/3 at myoblast contact sites, and optogenetic activation of protrusion in myofibers induced myoblast fusion. Genetic depletion of Arpc4 in myofibers markedly reduces membrane protrusions both *in vitro* and *in vivo*. These findings unveil a fundamental role for the Arp2/3 complex in postnatal muscle growth, and reveal a mechanism by which myofibers actively extend membrane protrusions to drive myoblast fusion. This repositions myofibers as active fusogenic partners and cell-autonomous drivers of their own nuclear accretion.

## INTRODUCTION

The actin nucleator Arp2/3 complex consists of two actin-related proteins (Arp2 and Arp3) and five additional subunits (ARPC1-5) ^1–3^. It promotes the formation of branched-actin networks by stimulating the growth of a new actin filament from the side of a pre-existing one ^4^. The Arp2/3 complex is ubiquitously expressed, and its genetic disruption *in vivo* is lethal in several organisms ^5–7^. The regulation of the actin cytoskeleton through the Arp2/3 complex is indispensable for a wide range of cellular processes, including cell migration, membrane trafficking, and the formation of cell-cell junctions ^8,9^.

Skeletal muscle accounts for approximately 40% of human body weight and is responsible for generating force and enabling body movement. Myofibers are large multinucleated muscle cells whose cytoplasm is mostly occupied with the contractile machinery ^10^. During embryonic development, precursor cells differentiate into myoblasts that fuse with each other to form multinucleated myotubes. Fusion of myoblasts depends on the formation of invasive protrusions, mediated by the Arp2/3 complex, to promote juxtaposition of cell membranes ^11–13^. After myoblast fusion, myotubes differentiate into myofibers, and myofiber nuclei (myonuclei) move to the periphery ^14^. We previously showed that peripheral myonuclei migration is a synchronized event occurring mainly at birth in mice, and that the Arp2/3 complex governs this process in an isoform-dependent manner ^15,16^. These findings raised the possibility that the Arp2/3 complex might also have a later, myofiber-intrinsic role in postnatal development, distinct from its earlier functions in myofiber formation and nuclear positioning. Postnatally, skeletal muscle growth relies on a pool of resident muscle stem cells, called the satellite cells (SCs), which localize between the basal lamina and the myofiber membrane ^17,18^. During the first 20 postnatal days, myonuclei number increases through the fusion of SCs-derived myoblasts with pre-existing myofibers, leading to an increase in myofiber size ^16,19,20^.

While the role of the Arp2/3 complex has been well-studied in myoblasts during embryogenesis, and its role in myonuclear positioning has been established, its specific function within myofibers during postnatal growth is entirely unknown. Here, we identify a myofiber-intrinsic, Arp2/3-dependent mechanism that actively drives myoblast fusion during postnatal development. Specifically, we show that myofibers form long-lived Arp2/3-enriched membrane protrusions at contact sites with myoblasts to promote fusion, and that loss of this activity impairs myonuclear accretion and muscle growth *in vivo*.

## RESULTS

### Loss of the Arp2/3 complex from myofibers during postnatal development compromises muscle function *in vivo*

The Arp2/3 complex is essential for embryonic skeletal muscle development ^11^. To study the role of the Arp2/3 complex in myofibers during postnatal skeletal muscle development we depleted Arpc4 (a core subunit of the Arp2/3 complex) specifically in myofibers using the HSA-MCM system ^21^. 4OH-tamoxifen was administered to both control Arpc4^fl/fl^ (Arpc4^WT^) and knockout HSA-MCM:Arpc4^fl/fl^ (Arpc4^-/-^) mice at post-natal (P) days 1-3 (Fig. 1A). At P7, Arpc4 protein level was reduced to half in the skeletal muscle of Arpc4^-/-^ mice, but not in other tissues (Fig. 1B,C and Sup. Fig. 1A). The expression of other Arp2/3 complex subunits (Arpc2, Arpc5 and Arpc5l), was also reduced in the skeletal muscle of Arpc4^-/-^ mice compared to Arpc4^WT^ (Fig. 1B and Sup. Fig. 1B-D).

**Figure 1.**
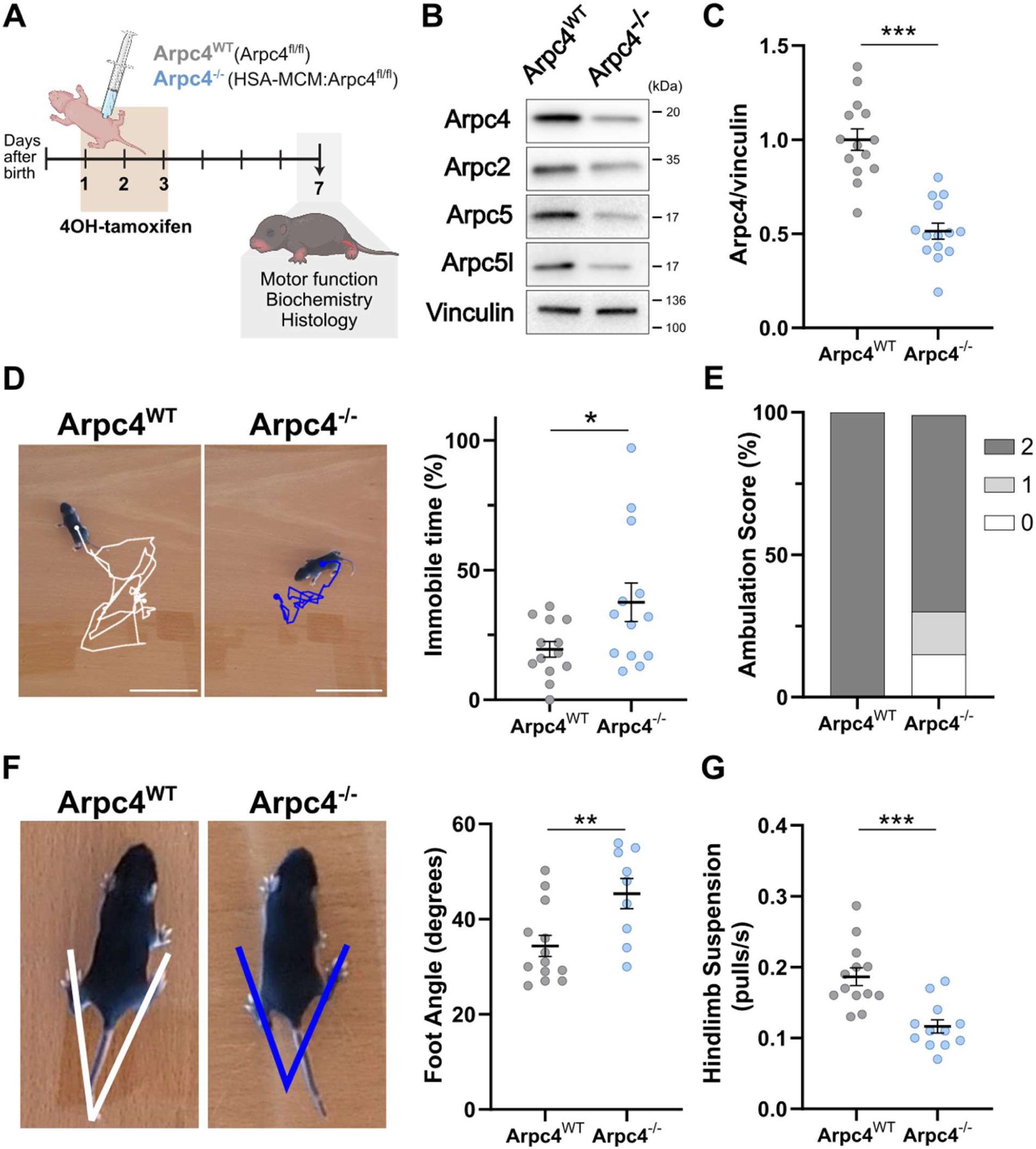
Depletion of Arp2/3 in myofibers leads to motor impairment during postnatal development. **A)** Schematic of the strategy used to generate skeletal muscle-specific Arpc4^-/-^mice. 4OH-tamoxifen administration at P1-P3 was used to induce skeletal muscle-specific Arpc4 depletion. At P7, locomotion was evaluated, and the hindlimb muscle was collected for analysis. **B)** Representative western blots (WB) for Arpc4, Arpc2, Arpc5, Arpc5l and Vinculin from skeletal muscle lysate of P7 Arpc4^WT^ and Arpc4^-/-^ mice. **C)** Quantification of Arpc4 protein levels by WB shown in (B), normalized against Vinculin. The results are shown as fold vs. the average of littermate Arpc4^WT^. **D)** (Left) Representative scheme of the trajectories of a P7 Arpc4^WT^ (white) and Arpc4^-/-^ (blue) mice during 3 min on an open field. Scale bar: 5 cm. (Right) Quantification of the percentage of time mice spent immobile in the open field. **E)** Quantification of ambulation score during open field observation (D). 0 = no movement; 1 = asymmetric movement; 2 = symmetric movement. **F)** (Left) Representative images of foot angles measured in P7 Arpc4^WT^ (white) and Arpc4^-/-^ (blue) mice while walking in a straight line, and respective quantification (right). **G)** Quantification of the number of pulls (body lifting) per second observed in mice while performing the hindlimb suspension test. Data are represented as mean ± s.e.m. in all expected I (n≥7 mice). *p≤0.05, **p≤0.01, ***p≤0.001 using unpaired 2-tailed Student’s t test or unpaired 2-tailed Mann-Whitney test, depending on normal distribution.

At P7, Arpc4^-/-^ mice were indistinguishable from Arpc4^WT^ animals and had a similar weight (Sup. Fig. 1E). Hematoxylin and Eosin (H&E) staining of gastrocnemius transverse sections showed that Arpc4^-/-^ skeletal muscle have an overall normal appearance, with no hallmarks of tissue damage, regeneration or immune cell infiltration (Sup. Fig. 1F). However, in the open field, Arpc4^⁻/⁻^ mice spent longer periods immobile compared to Arpc4^WT^ controls (Fig. 1D and Sup. Fig 1G). We then assessed the locomotor ability of the mice, using a set of tests appropriate for neonates ^22^. We observed that ∼30% of Arpc4^-/-^ mice either exhibited no exploration (movement not exceeding their own body length - ambulation score of 0), or presented asymmetric movements (mainly moving in circles - ambulation score 1) (Fig. 1E). In contrast, all Arpc4^WT^ controls had symmetric movements, being able to perform some straight and continuous locomotion (ambulation score 2) (Fig. 1E). Furthermore, for mice with symmetric locomotion, we observed that Arpc4^-/-^ mice foot angles were wider while walking, thus presenting splayed paws position (Fig. 1F). Finally, we performed the hindlimb suspension test and found that Arpc4^-/-^mice lifted their bodies (counted as pulls) with half the frequency of Arpc4^WT^ animals (Fig. 1G), suggesting decreased muscle strength. Our results indicate that the postnatal depletion of the Arp2/3 complex in myofibers leads to muscle weakness and abnormal locomotion.

### The Arp2/3 complex is involved in postnatal myofiber growth and nuclear accretion

To investigate how the depletion of the Arp2/3 complex in myofibers impacts mice locomotion, we performed histological analysis of the gastrocnemius of Arpc4^WT^ and Arpc4^-/-^ mice at P7. Immunostaining of the basal lamina showed that Arpc4^-/-^ myofibers do not have alterations in myofiber shape, since both circularity and roundness are similar to Arpc4^WT^ myofibers (Fig. 2A-C). However, when we analyzed the myofiber cross-sectional area (CSA), we found that the Arpc4^-/-^ muscle had a bigger percentage of smaller myofibers when compared to Arpc4^WT^ (Fig. 2A,D). Since postnatal myofiber growth largely depends on nuclear accretion *via* myoblast fusion ^19,20^, we quantified the number of myonuclei per myofiber. We observed that Arpc4^-/-^ myofibers have fewer myonuclei than Arpc4^WT^ myofibers, suggesting a decrease in myonuclear accretion (Fig. 2E,F). The peripheral myonuclei position, however, was not affected (Fig. 2E,G).

**Figure 2.**
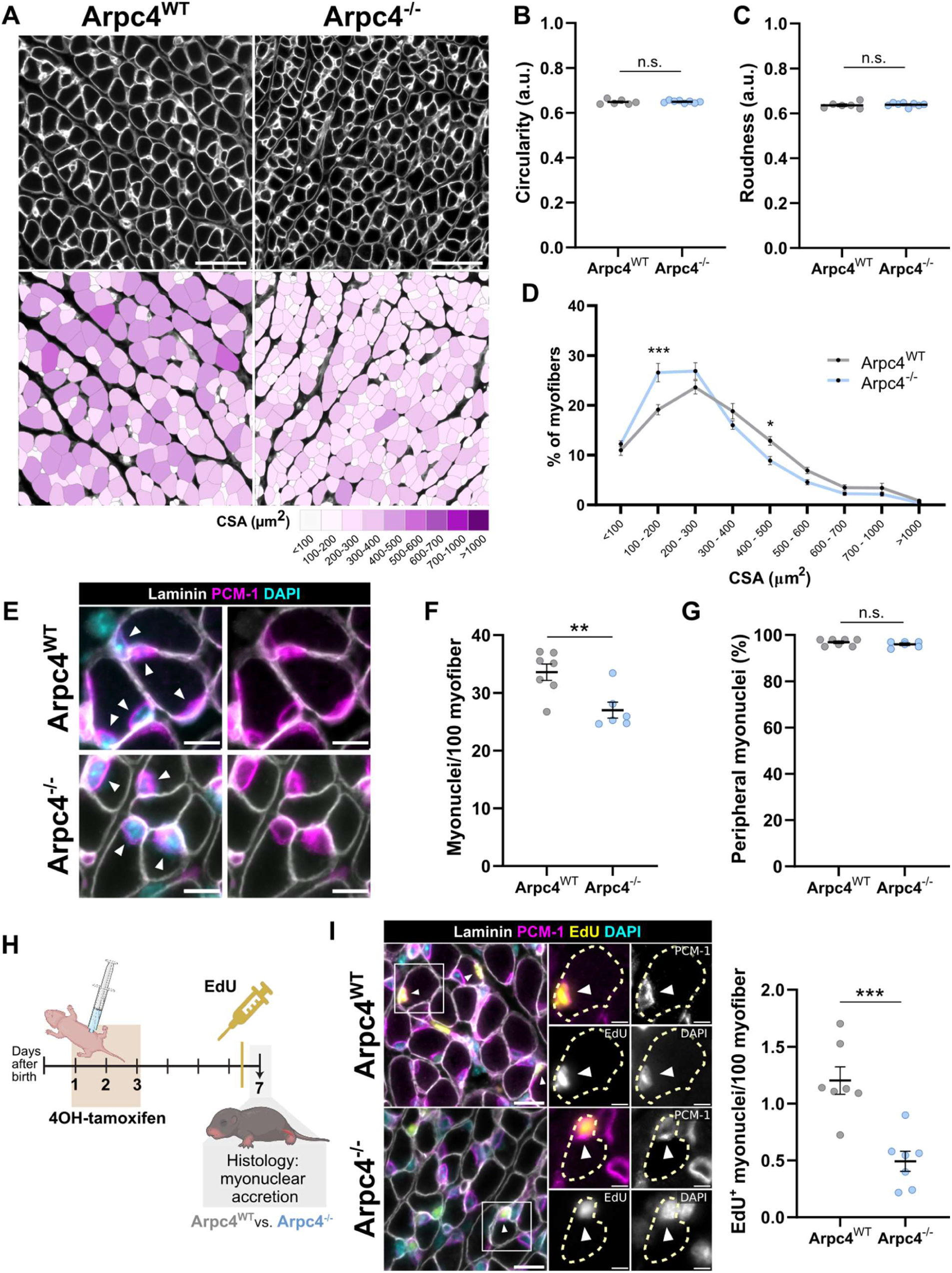
Arpc4^-/-^ mice have smaller myofibers and reduced myonuclear accretion. **A)** (Top) Representative images of the P7 Arpc4^WT^ and Arpc4^-/-^ mice gastrocnemius muscle cross-sections stained for the basal lamina (laminin, white). (Bottom) Top images, colour-coded by cross-sectional area (CSA), as indicated in the legend. Scale bar: 50 µm. **B-C)** Quantifications of the myofiber circularity (B) and roundness (C) of P7 Arpc4^WT^ and Arpc4^-/-^ mice myofibers. **D)** Frequency distribution of myofiber CSA of P7 Arpc4^WT^ and Arpc4^-/-^ mice. **E)** Representative images of cross-sectioned hindlimb muscle from P7 Arpc4^WT^ and Arpc4^-/-^ mice, stained for the basal lamina (laminin, white), myonuclei (PCM-1, magenta) and DNA (DAPI, cyan). White arrows identify myonuclei. Scale bar: 10 µm. **F-G)** Quantification of myonuclei number per 100 myofiber (F) and percentage of myonuclei localized at the periphery of the myofibers (G), in the cross-sections as in (E). **H)** Schematic of the strategy used to evaluate cell fusion and myonuclear accretion in Arpc4^WT^ and Arpc4^-/-^ mice. 4OH-tamoxifen administration at P1-P3 was used to induce skeletal muscle-specific Arpc4 depletion. EdU was administered at 18h before mice were sacrificed, and the gastrocnemius muscle was collected at P7. **I)** (Left) Representative images of cross-sectioned muscle from P7 Arpc4^WT^ and Arpc4^-/-^ mice, stained for the basal lamina (laminin, white), myonuclei (PCM-1, magenta), EdU (yellow) and DNA (DAPI, cyan). White arrows identify EdU^+^ myonuclei. Scale bar: 15 µm. Magnifications corresponding to the white squares are shown on the right of each image, with each channel shown separately. The yellow dashed line shows the myofiber delimitation through laminin staining. Scale bar: 5 µm. (Right) Quantification of the number of EdU^+^ myonuclei per 100 myofibers. Data are represented as mean ± s.e.m. (n≥6 mice). *p≤0.05, **p≤0.01, ***p≤0.001 using unpaired 2-tailed Student’s t test in all except (D), in which two-way ANOVA with Sidak’s post-hoc test was used.

In higher eukaryotes, three Arp2/3 subunits (Arp3, Arpc1 and Arpc5) are encoded by two separate genes, leading to distinct isoforms of the same subunit ^23–25^. This means that eight different Arp2/3 complexes, with distinct properties, may be generated within the same cell ^26–28^. We previously showed that the Arp2/3 complex has distinct functions in skeletal muscle development, depending on the Arpc5 isoform ^15,29^. To understand if the alterations in Arpc4^-/-^mice were Arpc5-isoform-dependent, we specifically depleted Arpc5 or Arpc5l from the myofibers during postnatal development, using the HSA-MCM system. We confirmed that both Arpc5 and Arpc5l are specifically depleted from the skeletal muscle of Arpc5^-/-^ and Arpc5l^-/-^ mice, respectively (Sup. Fig. 2A-F). We found no changes in myofiber CSA, morphology, or myonuclear number in either Arpc5^-/-^ or Arpc5l^-/-^ mice when compared with respective controls (Sup. Fig. 2G–N). Therefore, the reduction in myofiber CSA and myonuclei number observed in Arpc4^⁻/⁻^ mice is independent of the Arpc5 isoforms.

We next tested whether the decrease in CSA and myonuclei number in Arpc4^-/-^ myofibers was due to reduced myoblast-myofiber fusion. To this end, we administered EdU 18h before muscle collection in both Arpc4^WT^ and Arpc4^-/-^ mice, which will label proliferating cells (including activated SCs and myoblasts) (Fig. 2H). Since myofibers are post-mitotic, myonuclei are not labelled with EdU. Thus, all EdU+ myonuclei correspond to newly added nuclei to the myofibers through myoblast fusion ^16,30^. We observed that myonuclear accretion is strongly reduced in Arpc4^-/-^myofibers (Fig. 2I), revealing that the Arp2/3 complex in myofibers is required for their fusion with myoblasts.

### Depletion of Arpc4 in myofibers prevents fusion of activated SCs

During postnatal development, SCs become activated, proliferate and differentiate into myoblasts committed to fuse with myofibers ^31,32^. We investigated whether SCs activation was impaired upon depletion of Arp2/3 in myofibers. We used Pax7 as a marker for the SCs, together with its localization below the basal lamina. The proliferative marker Ki-67 was used to distinguish quiescent from proliferative activated SCs, and the myogenic factor MyoD was used to identify committed myoblasts (Fig. 3A-C). Our analysis showed that the number of quiescent SCs (Pax7^+^/Ki-67^-^) in Arpc4^-/-^ mice remained unaltered, indicating that SCs pool is not affected (Fig. 3A). However, there was an increase in the number of activated SCs (Pax7^+^/Ki-67^+^), as well as committed myoblasts (MyoD^+^), between the basal lamina and the plasma membrane of Arpc4^-/-^myofibers (Fig. 3A,B). When we analysed longitudinal sections, we observed that Pax7^+^/Ki-67^+^ cells accumulated in clusters in close proximity to the myofibers, confirming that activated SCs failed to fuse with Arpc4-depleted myofibers (Fig. 3C).

**Figure 3.**
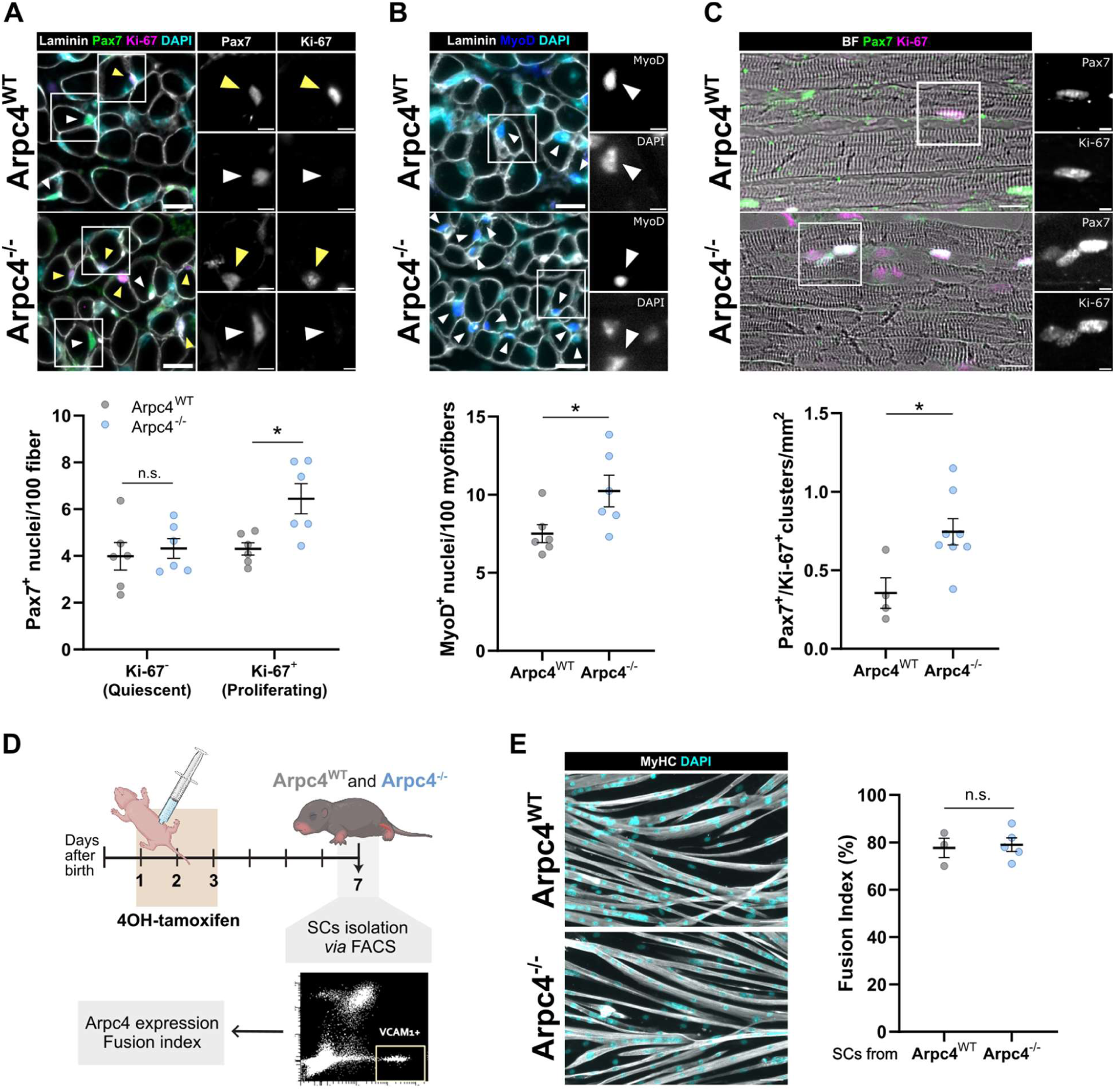
Activated SCs accumulate around myofibers in Arpc4^-/-^ muscle. **A)** (Top) Representative images of cross-sectioned gastrocnemius muscle from P7 Arpc4^WT^ and Arpc4^-/-^mice, stained for the basal lamina (laminin, white), Pax7 (green), proliferation (Ki-67, magenta) and DNA (DAPI, cyan). White arrows identify quiescent SCs (Pax7^+^/Ki-67^-^), while yellow arrows identify proliferating SCs (Pax7^+^/Ki-67^+^). Scale bar: 15 µm. Magnifications corresponding to the white squares are shown on the right of each image, with Pax7 and Ki-67 signals shown separately. Scale bar: 5 µm. (Bottom) Quantification of the number of quiescent and proliferating SCs per 100 myofibers. **B)** (Top) Representative images of cross-sectioned gastrocnemius muscle from P7 Arpc4^WT^ and Arpc4^-/-^ mice, stained for the basal lamina (laminin, white), MyoD (blue) and DNA (DAPI, cyan). White arrows identify MyoD^+^ nuclei. Scale bar: 15 µm. Magnifications corresponding to the white squares are shown on the right of each image, with MyoD and DAPI shown separately. Scale bar: 5 µm. (Bottom) Quantification of the number of MyoD^+^ nuclei per 100 myofibers. **C)** (Top) Representative images of longitudinal gastrocnemius sections from P7 Arpc4^WT^ and Arpc4^-/-^ mice, stained Pax7 (green) and proliferation (Ki-67, magenta). Myofibers are shown in the brightfield (BF). Scale bar: 15 µm. Magnifications corresponding to the white squares are shown on the right of each image, with Pax7 and Ki-67 shown separately. Clusters of SCs were counted when two or more Pax7^+^/Ki-67^+^ nuclei were in a region smaller than 30 µm x 12 µm. Scale bar: 5 µm. (Bottom) quantification of the number of Pax7^+^/Ki-67^+^ clusters per mm^2^. **D)** Schematic representation of the strategy used to isolate a pure population of SCs from P7 Arpc4^WT^ and Arpc4^-/-^ skeletal muscle by FACS-sorting. **E)** (Left) Representative images of myofibers generated from SCs isolated from P7 Arpc4^WT^ and Arpc4^-/-^mice, stained for Myosin Heavy Chain (MyHC, white) and DNA (DAPI, cyan), and (Right) the respective fusion index. Data are represented as mean ± s.e.m. (n≥3 mice). *p≤0.05, **p≤0.01, ***p≤0.001 using unpaired 2-tailed Student’s t test or unpaired 2-tailed Mann-Whitney test, depending on normal distribution, in all except (D), in which two-way ANOVA with Sidak’s post-hoc test was used.

To ensure that the ability of SCs to fuse was unaffected in Arpc4^-/-^ mice, we FACS-sorted SCs from the muscle of P7 Arpc4^WT^ and Arpc4^-/-^ mice and assessed their differentiation *in vitro.* (Fig. 3D). These SCs were positive for myogenic markers, including Pax7 and Desmin, and expressed Arpc4 at the same protein levels, as expected (Sup. Fig. 3A,B). Upon differentiation, no differences were observed in the fusion index of SCs isolated from Arpc4^WT^ or Arpc4^-/-^ mice (Fig. 3E), confirming that the intrinsic ability of SCs to fuse was not affected by myofiber-specific depletion of Arpc4 *in vivo*. Altogether, our results demonstrate that the loss of Arpc4 in myofibers impairs fusion with functionally activated myoblasts, thus highlighting the role of the Arp2/3 complex in myofibers for myonuclear accretion and muscle growth.

### Myofibers generate Arp2/3-enriched membrane protrusions involved in myofiber fusion

To further investigate the role of the Arp2/3 complex in myofibers during myonuclear accretion, we developed a co-culture system to visualize myoblast-myofiber interaction and fusion *in vitro*. This system enables us to investigate myofiber-specific mechanisms that are involved in the fusion process. Briefly, we differentiated FACS-sorted SCs into myofibers over 3 days (D3), and co-cultured them with mononucleated myoblasts (Fig. 4A). To observe fusion, we pre-labeled the myoblasts with EdU to follow their accretion into pre-existing myofibers (Fig. 4B). After 48h of co-culture, we detected EdU^+^ nuclei in pre-existing myofibers (Fig. 4C), thus indicating fusion of myoblasts into myofibers, and validating this system to study myofiber myonuclear accretion.

**Figure 4.**
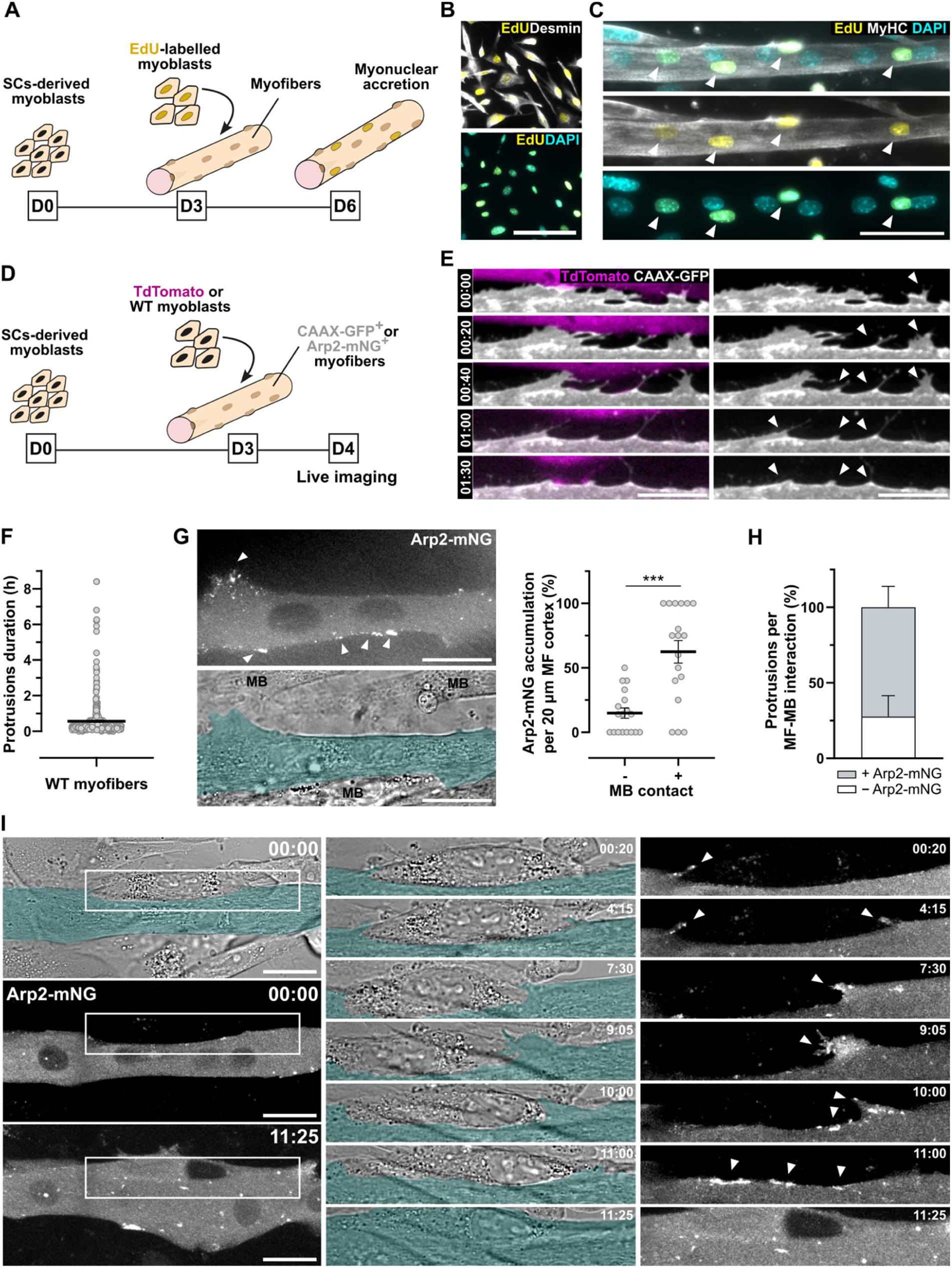
Myofibers create long-lived membrane protrusions and accumulate Arp2/3 at contact sites with myoblasts. **A)** Schematic of the co-culture system development to study myoblast-myofiber fusion *in vitro*. D3 myofibers were co-cultured with EdU-labeled SCs-derived myoblasts. Myofibers were analyzed 2 days after co-culture. **B)** Representative images of WT myoblasts after 24h incubation with EdU, stained for Desmin (white), EdU (yellow) and DAPI (cyan). Scale bar: 100 µm. **C)** Representative images of the co-culture between D3 myotubes and EdU-labeled SCs in (B), stained for MyHC (white), EdU (yellow) and DAPI (cyan). White arrows identify EdU^+^ myonuclei fused in pre-existing myofibers (with more than 3 EdU^-^ nuclei). Scale bar: 50 µm. **D)** Schematic of the co-culture system development for live imaging study of myoblast-myofiber interaction and fusion *in vitro*. D3 myofibers expressing CAAX-GFP or Arp2-mNG were co-cultured with tdTomato or unlabeled myoblasts, respectively. **E)** Frames from a time-lapse video showing that myofibers create long-lived membrane protrusions (white arrows) upon interaction with myoblasts (magenta). Myofiber membrane is labeled with a CAAX-GFP (white). Time in hh:min; Scale bar: 20 µm. **F)** Quantification of the duration of myofiber protrusions created during interaction with myoblasts, as in (E). Data are represented as mean ± s.e.m. (n=436 protrusions, from 3 independent experiments). **G)** (Left) Representative image of the localization of Arp2-mNG (white) in the myofibers (pseudocoloured in cyan). White arrows identify accumulation of Arp2-mNG in regions of contact with myoblasts (MB). Scale bar: 20 µm. (Right) Quantification of the percentage of Arp2-mNG accumulation in regions of the myofiber (MF) with or without a neighbouring myoblast (MB). Data are represented as mean ± s.e.m. (n=18 myofibers, from 3 independent experiments). *p≤0.05, **p≤0.01, ***p≤0.001 using unpaired 2-tailed Mann-Whitney test. **H)** Quantification of the percentage of myofiber protrusions that have or do not Arp2-mNG accumulation. **I)** Frames from a time-lapse video of Arp2-mNG (white) in myofibers (pseudocoloured in cyan), during interaction with unlabeled myoblasts. In the BF, it is possible to follow the position of the unlabeled myoblast relative to the myofiber (pseudocoloured in cyan). Fusion is observed at 11:25 (hh:min). Magnifications corresponding to the white rectangles in different moments are shown at the center (BF) and at the right (Arp2-mNG). Scale bar: 20 µm.

We examined the interaction between myofibers and myoblasts *in vitro* by performing live imaging of tdTomato-myoblasts co-cultured with myofibers expressing a membrane-targeted GFP (CAAX-GFP) (Fig. 4D). We found that myofibers form numerous membrane protrusions when interacting with myoblasts (Fig. 4E and Sup. Movie 1). These protrusions are long-lived, with an average lifetime of 30 min, and some persist for longer than 2h (Fig. 4E,F). Given the known role of the Arp2/3 complex in the formation of membrane protrusions, we next explored its dynamics in myofibers during interaction with myoblasts ^9^. We performed live imaging of unlabeled myoblasts co-cultured with myofibers expressing Arp2-mNG (Fig. 4D). We found that Arp2-mNG accumulates 4 times more in regions where the myofiber is in contact with myoblasts, when compared to regions without any contact (Fig. 4G). In addition, we observed that Arp2-mNG accumulates in 70% of myofiber protrusions in contact with myoblasts (Fig. 4H), suggesting a role for the Arp2/3 complex in the formation and maintenance of such structures. To understand if the myofiber protrusions were involved in their fusion with myoblasts, we performed time-lapse imaging to visualize fusion events between Arp2-mNG^+^ myofibers and unlabeled myoblasts (Fig. 4I and Sup. Movie 2). We observed the formation of myofiber protrusions containing Arp2-mNG, followed by further accumulation of Arp2-mNG in the myofiber-myoblast contact interface. Finally, these dynamic interactions culminated in the fusion between the myofiber and the myoblast, suggesting that the Arp2/3-containing myofiber protrusions are involved in the fusion (Fig. 4I).

To test if membrane protrusions were sufficient to promote fusion of myofibers, we induced Arp2/3-dependent membrane protrusion through optogenetic activation of Cdc42 in myofibers co-cultured with unlabeled myoblast (Fig. 5A) ^33^. Illumination with 488 nm light triggered extensive membrane protrusion formation in mCherry-ITSN-CRY2 (Cdc42^Opto^) but not mCherry-CRY2 (mCherry^Opto^) myofibers (Fig. 5B). Strikingly, 50% of Cdc42^Opto^ myofibers fused to myoblasts during 12 h optogenetic activation, whereas no fusion was observed in mCherry^Opto^ myofibers during the same time interval (Fig. 5B and Sup Movie 3-4). Therefore, activation of Arp2/3-dependent membrane protrusion in myofibers is sufficient to induce their fusion with myoblasts.

**Figure 5.**
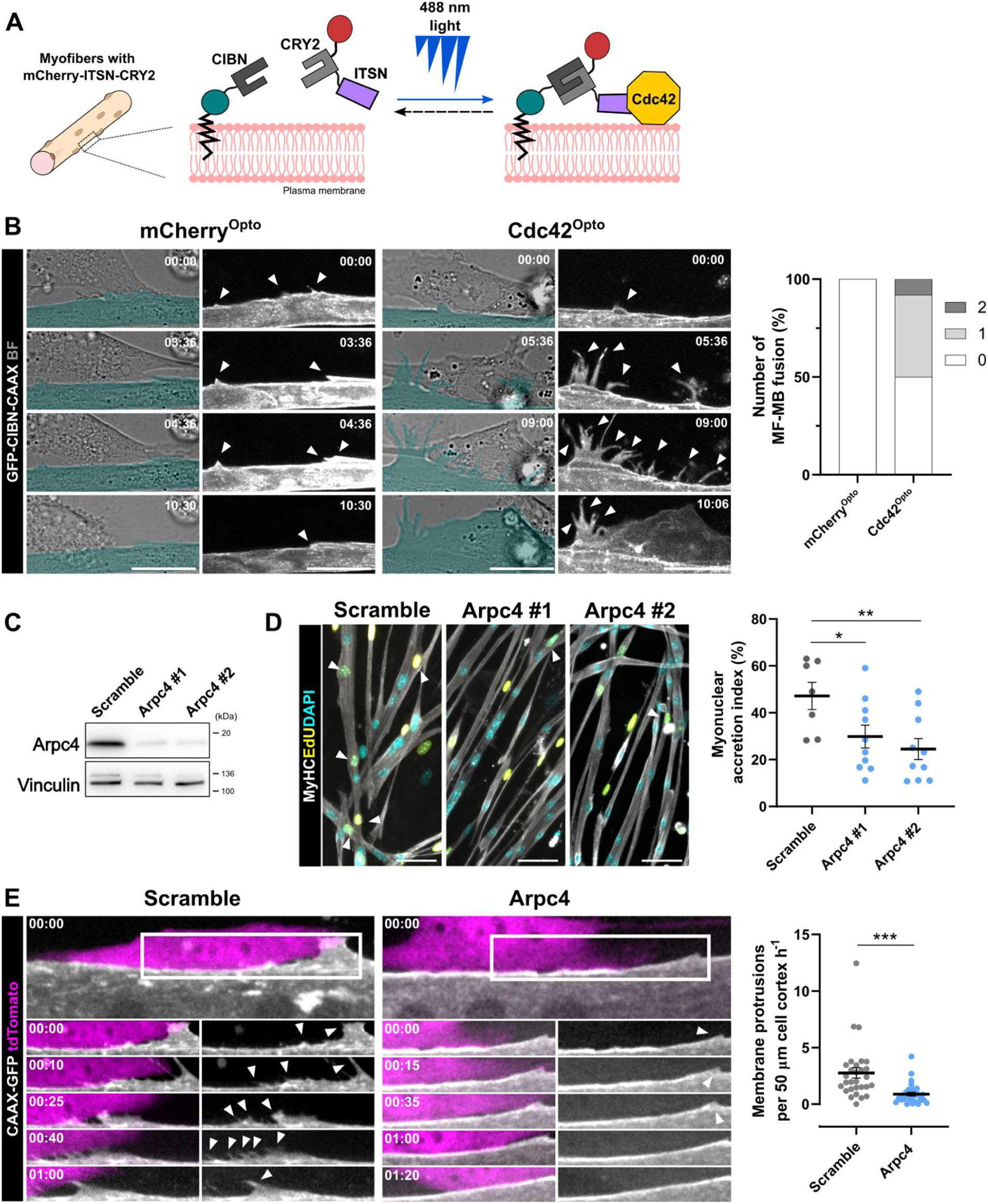
Myofibers create Arp2/3-dependent membrane protrusions required for fusion with myoblasts. **A)** Schematic representation of the optogenetic tool used to activate Cdc42 in myofibers. **B)** (Left) Frames from a time-lapse video showing the interaction between unlabeled myoblasts and myofibers expressing GFP-CIBN-CAAX (white), and mCherry-CRY2 (mCherry^Opto^) or mCherry-ITSN-CRY2 (Cdc42^Opto^). In BF, the myofiber is pseudocoloured in cyan. Time in hh:min; Scale bar: 20 µm. (Right) Quantification of the percentage of myofibers that fused with 0, 1 or 2 myoblasts during 12h of optogenetic activation. **C)** Western blots for Arpc4 and Vinculin in D3 myotubes treated with Arpc4 #1, Arpc4 #2 or Scramble siRNA. **D)** (Left) Representative images of the co-culture between D3 WT or Arpc4-depleted myotubes, and EdU-labeled myoblasts, stained for MyHC (white), EdU (yellow) and DAPI (cyan). White arrows identify EdU+ myonuclei in pre-existing myofibers (with more than 3 EdU^-^ myonuclei). Scale bar: 50 µm. (Right) Quantification of the percentage of EdU^+^ nuclei that fused with pre-existing myofibers (myonuclear accretion index). **E)** (Left) Frames from a time-lapse video showing the interaction between tdTomato myoblasts (magenta) and CAAX-GFP (white) myofibers (scramble and Arpc4 siRNA #2). The white rectangle represents the 50 µm of the myofiber membrane that was analyzed. Time in hh:min; Scale bar: 25 µm. White arrows identify membrane protrusions in the myofibers. (Right) Quantification of the number of membrane protrusions formed per hour in 50 µm of the myofiber membrane in contact with tdTomato-myoblasts. Data are represented as mean ± s.e.m. (n≥29 myofiber-myoblast interaction, from 3 independent experiments). *p≤0.05, **p≤0.01, ***p≤0.001 using unpaired 2-tailed Mann-Whitney test.

### The Arp2/3 complex is required for the formation of myofiber protrusions during postnatal development

To understand if the myofiber protrusions are dependent on the Arp2/3 complex, we investigated the interaction between myofibers and myoblasts after Arpc4 depletion. We differentiated FACS-sorted SCs into myofibers depleted of Arpc4, either by siRNA or by Cre-Recombinase expression in Arpc4^fl/fl^ cells (Fig. 5C and Sup. Fig. 4A). We co-cultured those myofibers with EdU^+^ myoblasts, and quantified the percentage of myoblasts that fused into pre-existing myofibers, defined as Myonuclear Accretion Index (MAI). We observed a significant decrease in the MAI when WT myoblasts were co-cultured with Arpc4-depleted myofibers, either by siRNA or by Cre-recombinase expression (Fig. 5D and Sup. Fig. 4B), as observed *in vivo*. We then performed live imaging of tdTomato-myoblasts co-cultured with CAAX-GFP myofibers to analyze the formation of myofiber protrusions upon depletion of Arpc4. We found that Arpc4-depleted myofibers exhibit three times fewer protrusions per hour during interaction with myoblasts, when compared to control cells (Fig. 5E and Sup. Movie 5,6). Our results demonstrate that the Arp2/3 complex in myofibers is required for the formation of membrane protrusions, which are sufficient to induce myofibers fusion with myoblasts.

To assess whether the Arp2/3-dependent myofiber protrusions are relevant for postnatal myonuclear accretion, we analyzed cross-sections of gastrocnemius muscle of P7 Arpc4^WT^ and Arpc4^-/-^ mice by TEM (Fig. 6A). Strikingly, we found that Arpc4^WT^ myofibers create protrusions in ∼80% of the interactions with myoblasts, in contrast to Arpc4^-/-^ where 60% of myofibers do not create any membrane protrusion (Fig. 6B). Quantification of the number of membrane protrusions revealed that Arpc4^-/-^ myofibers exhibited a strong decrease when compared to Arpc4^WT^ (Fig. 6C). Furthermore, the few protrusions generated by Arpc4^-/-^ myofibers were smaller than the ones formed in Arpc4^WT^ mice (Fig. 6D). Taken together, our data demonstrate that the Arp2/3 complex is required for the formation of myofiber protrusions that mediate the interaction and fusion with myoblasts during postnatal development.

**Figure 6.**
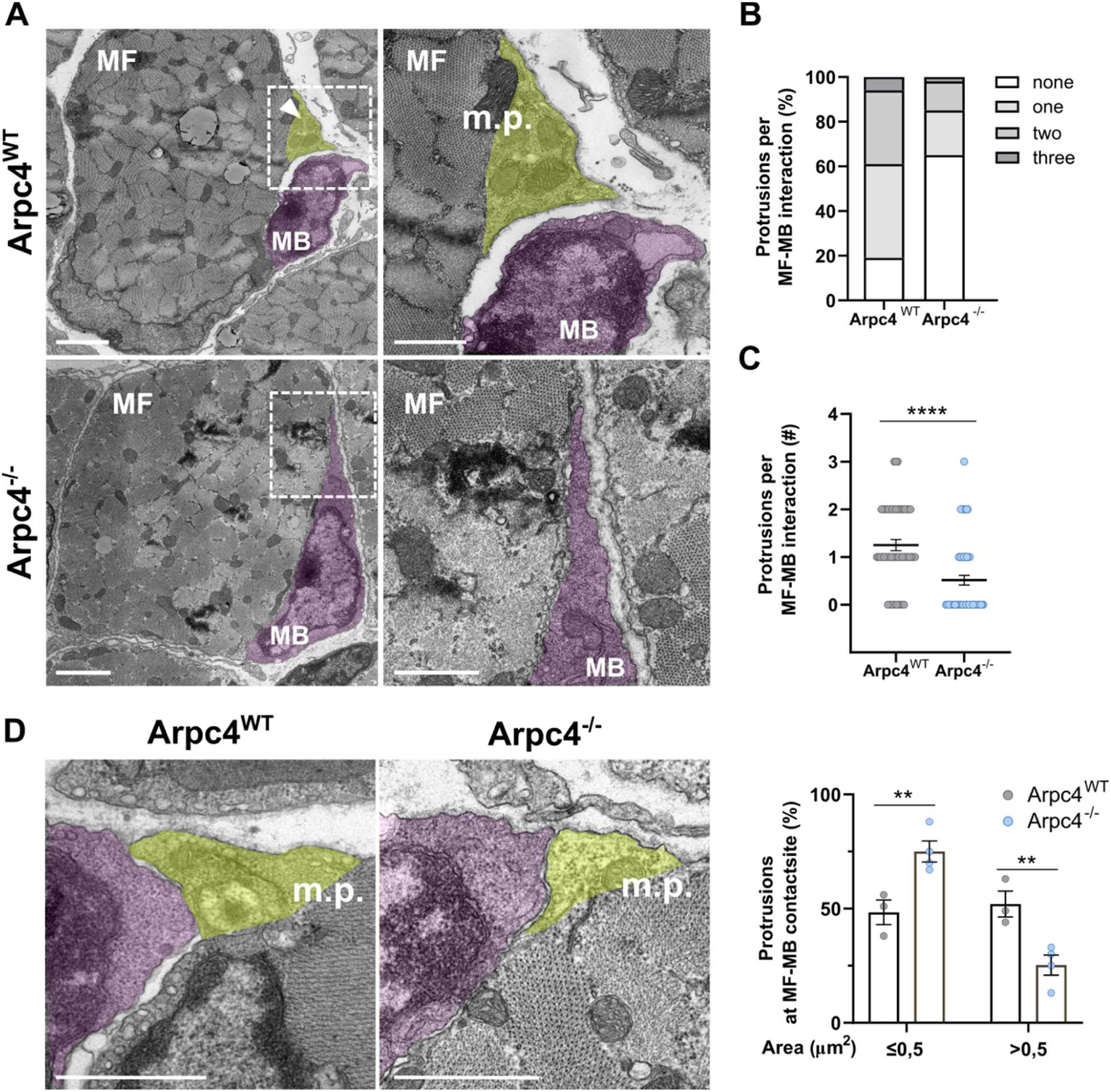
Arp2/3-dependent membrane protrusions are involved in myofibers-myoblast interaction during postnatal development. **A)** TEM representative images from P7 Arpc4^WT^ and Arpc4^-/-^ mice gastrocnemius cross-sections. Note a myofiber (MF) membrane protrusion (m.p.) towards a myoblast (MB, pseudocoloured in magenta) in Arpc4^WT^ but not in Arpc4^-/-^. Scale bar: 2µm. Magnifications corresponding to the white square are shown on the right of each image. Scale bar: 1µm. **B)** Percentage of myofibers with 0, 1, 2 or 3 membrane protrusions per MF-MB contact site in Arpc4^WT^ and Arpc4^-/-^ mice. **C)** Quantification of the number of myofiber protrusions per MF-MB contact site in Arpc4^WT^ and Arpc4^-/-^ mice. **D) (Left)** TEM representative images highlighting myofiber protrusions (pseudocoloured in yellow). Scale bar: 1µm. (Right) Quantification of the percentage of myofiber protrusions with less or more than 0.5 μm^2^ of cross-sectional area in Arpc4^WT^ with Arpc4^-/-^ mice. Data is represented as mean ± s.e.m. *p≤0.05, **p≤0.01, ***p≤0.001 using two-way ANOVA with Sidak’s post-hoc test.

## DISCUSSION

Postnatal muscle growth relies on the fusion of myoblasts with pre-existing myofibers, resulting in a progressive increase in myofiber size and myonuclei number. The regulation of this process has been mostly attributed to the fusing cell, the myoblast ^34^. Here, we demonstrate for the first time that the receiving cell, the mature myofiber, is also an active regulator of the fusion process. We showed that myofibers extend long-lived Arp2/3-enriched membrane protrusions towards neighbouring SCs, promoting their fusion. In the absence of the Arp2/3 complex, the myofiber membrane is less dynamic during cell-cell interaction and fusion with myoblasts is impaired. Additionally, we showed that postnatal depletion of the Arp2/3 complex in myofibers impairs SCs-dependent myofiber growth, resulting in gait abnormalities and muscle weakness.

Prior studies established Arp2/3-dependent invasive protrusions in myoblast fusion during embryogenesis and regeneration ^11,35,36^. Our work revealed that myofiber-specific depletion of the Arp2/3 complex during postnatal development leads to smaller myofibers with fewer myonuclei, without altering SC activation or intrinsic fusion competence. EdU labeling of proliferating SCs *in vivo* revealed that fusion of myoblasts to Arp2/3-depleted myofibers was strongly compromised, leading to an accumulation of activated SCs in the skeletal muscle of Arpc4^-/-^ mice. Our findings reveal that myofibers are active players in regulating myonuclear accretion through Arp2/3-dependent mechanisms.

We previously showed that the Arp2/3 complex containing Arpc5l isoform is required for the migration of myonuclei to the periphery of myofibers, through a mechanism involving the organization of desmin ^15^. We also showed that most myonuclei migrate to the periphery before P1, and postnatal myoblast fusion does not result in centrally nucleated myofibers as observed in the embryo ^16^. Our results demonstrate that the Arp2/3 complex is not required for anchoring myonuclei at the periphery, since postnatal depletion of the Arp2/3 complex in myofibers did not affect myonuclear positioning. Importantly, we found that the Arp2/3 complex acts as an essential regulator of myofiber fusion during postnatal development, independently of the Arpc5 isoforms. This is particularly noteworthy given that Arpc5 isoforms govern distinct Arp2/3 functions in muscle ^15,29^, suggesting that the postnatal fusion machinery relies on a different mechanism that is robust to isoform variation.

Our work revealed that myofibers form long-lived Arp2/3-dependent membrane protrusions when interacting with neighbouring myoblasts, which promote myofiber fusion. Strikingly, optogenetic activation of Cdc42-driven protrusions in myofibers was sufficient to induce fusion with adjacent myoblasts in the absence of any other stimulus, establishing a direct causal link between myofiber protrusions and the fusion event. This gain-of-function result argues that the protrusive activity of the myofiber is not merely permissive but is itself a driver of myonuclear accretion. Several specialised membrane structures have been identified as key contributors to cell-cell communication and adhesion, especially during development ^37–40^. In the epithelia, cells form Arp2/3-dependent protrusions enriched in adhesion molecules (E-cadherin) during the formation of epithelial sheets, increasing the efficiency of cell-cell adhesion ^40,41^. We hypothesize that a similar mechanism may be involved in myofiber fusion, in which myofibers create Arp2/3-dependent membrane protrusions to facilitate the interaction with myoblasts and potentiate cell adhesion for fusion.

In Dr*osophila* embryo, trans-interaction between cell-type-specific adhesion molecules triggers Arp2/3-dependent actin cytoskeleton rearrangement, bringing membranes into juxtaposition and promoting myoblast fusion ^13,42,43^. We also observed an accumulation of the Arp2/3 complex in regions of the myofiber in proximity to myoblasts, which may be recruited by unidentified adhesion molecules in mammalian myofibers ^44^.

Our work demonstrates that myofibers have Arp2/3-dependent mechanisms to actively regulate myofiber growth. In the absence of the Arp2/3 complex in myofibers, neonate mice exhibit increased foot angles while walking and muscle weakness when performing the hindlimb suspension test. Muscle weakness, impaired ambulation and smaller myofibers are hallmarks of some congenital myopathies, namely centronuclear myopathies (CNM) ^45^. Additional work should address whether the function of the Arp2/3 complex is altered in disease and may contribute to its pathophysiology.

Collectively, our study unveils a fundamental cell-autonomous role for the Arp2/3 complex in postnatal muscle growth. We demonstrate that myofibers are not passive recipients, but active fusogenic partners that generate Arp2/3-dependent membrane protrusions, which are necessary for myonuclear accretion *in vivo* and sufficient to drive fusion upon activation. These findings reposition myofibers as drivers of their own nuclear accretion.

## Acknowledgments

We are grateful to all members of the Edgar Gomes, Michael Way and Carolyn Moore Laboratories for their continued support, scientific discussions and helpful comments on this work. We specifically thank: Sara Ferreira for her contribution with mouse samples; Dr. Helena Pinheiro for her help setting up the behaviour tests in the neonates; Dr. Ana Raquel Pereira for her help in generating important results through optogenetic experiments on Cdc42 activation in myofibers. We additionally thank the Neves & Victor Lab at the Gulbenkian Institute for Molecular Medicine (GIMM) for kindly providing the Pax7^iCreERT2;ROSA26LSL^-tdTomato mouse model. We also extend a special thank you to Elizabeth H. Chen for providing us with the Arp2-mNG construct, and to Bruno Cadot for his assistance with the CSA analysis. We finally thank the technical support of the facilities at GIMM, namely the Rodent, the Comparative Pathology, the Flow Cytometry and the Bioimaging (funded by PPBI-POCI-01-0145-FEDER-022122), as well as the Molecular and Transgenic Tools Platform (MTTP) of Champalimaud Foundation for cloning and AAV production.

## Author contributions

C.S. and S.D.F carried out experiments and analyzed the data, with contributions from G.M. N.K., who generated the Arpc5 and Arpc5l mice, and G.M. performed the experiments on those strains. C.S., S.D.F. and G.M. prepared the figures. C.S., S.D.F. and E.R.G. designed experiments and wrote the manuscript. E.R.G and M.W. supervised the project and secured funding. All the authors participated in the critical review and revision of the manuscript.

## Data and materials availability

All data are available in the main text or the supplementary materials.

## MATERIAL AND METHODS

### Mice

Skeletal muscle-specific Arp2/3 knockout mice strains were generated by breeding mice floxed for Arpc4 (Arpc4^-/-^) ^46^, Arpc5 (Arpc5^-/-^) or Arpc5l (Arpc5l^-/-^), with mice expressing MerCreMer double fusion protein under the human ACTA1 promoter (HSA-MCM) (Jackson Laboratory). Cre-negative littermates were used as controls. Both males and females were used without distinction. 4-Hydroxytamoxifen (4OH-Tamoxifen, Sigma, H6278) was injected intraperitoneally (50 µl of 1 mg/mL solution) at P1, P2 and P3. At P7-P8, motor function was evaluated and/or animals were euthanized by decapitation. Pax7^iCreERT2;ROSA26LSL^-tdTomato mouse model (Pawlikowski et al., 2015), kindly provided by Neves & Sousa-Victor Lab (GIMM, Lisbon, Portugal) were used to isolate tdTomato SCs. Three daily consecutive tamoxifen (Sigma-Aldrich, T5648) intraperitoneal injections (∼75 mg/kg) in mice younger than 8 weeks were used to permanently label Pax7-positive SCs with tdTomato, and SCs were isolated *via* FACS-sorting.

All mice were housed in a room maintained with a 12-12 hour light-dark cycle and received food and water ad libitum at the Rodent Platform at the Gulbenkian Institute for Molecular Medicine. All the procedures were approved by the Institutional Ethics Committee and followed the guidelines of the Portuguese National Research Council Guide (DGAV Project Licenses n° 022871/2016 and 004560/2022) for the care and use of laboratory animals.

### Motor function evaluation

To evaluate motor function in pups we used a group of tests appropriate to neonates, namely the ambulation score and angles, and the hindlimb suspension test ^22^. Tests were performed early in the morning to exclude differences in behaviour due to the time of day. In the ambulation score, pups were placed in a clear enclosure for 3 minutes, and the ambulation was scored according to the defined criteria (0 = if movement not exceeding their own body length; 1 = crawling/walking with asymmetric limb movement, when mainly moving in circles; 2 = crawling/walking with symmetric limb movement, in straight lines). Using the video recordings, we tracked the animals using ManualTrack on ImageJ, and measured the immobile time, considering immobile when the mice spent longer than 5 sec without moving. We also measured the pup’s foot angles by drawing a line from the heel to the tip of the longest (middle) toe. Foot angles were measured only when the pup was performing a full stride in a straight line, and both feet were flat on the ground. In the hindlimb suspension test, pups were placed gently, facing down, into a 50 ml conical tube with their hind legs hanging over the rim. We counted how many times the animals lifted their bodies using the hindlimbs (pulls). This was repeated 3 times, with 30 sec of resting time between each trial. The final result is the average of the 3 repetitions.

### SCs isolation and cell culture

SCs were isolated from limb skeletal muscles of Arpc4^WT^ or Pax7-tdTomato mice, younger than 8 weeks of age, as previously described^47^. Briefly, muscle was mechanically minced and digested for 1 h using NB4 collagenase (12 mg/ml) (SERVA) and Dispase II (100 U/ml) (Roche) at 37°C under agitation, and with TrypLE Express (Gibco) for 5 min. After filtering and centrifuging, Arpc4 cells were stained with PE-conjugated anti-Sca1, -CD31, and -CD45 antibodies and an anti-VCAM1 primary antibody, followed by a secondary AlexaFluor 488-conjugated secondary antibody (Table 1). Sca1^-^/CD31^-^/CD45^-^ and VCAM1^+^ cells or tdTomato cells were selected either with a BD FACSAria Fusion or a BD FACSAria III Cell Sorter (BD Biosciences). Live cells were gated using Propidium Iodide or Live/Dead staining kit (Life Technologies). Cell sorting was performed at the Flow Cytometry Platform of the Gulbenkian Institute for Molecular Medicine. Cells were cultured in laminin-coated dishes (3 µg/cm2) (Millipore) in DMEM-F12 (Gibco, 31331028) medium with 15% FBS and 1% PenStrep, and supplemented with bFGF (2.5 µg/ml, Sigma Aldrich) and B27 supplement without vitamin A (Gibco). Cells were maintained at 37°C and 5% CO2. For passage, cells were washed with DPBS and detached with TrypLE Express (Gibco) at 37 °C for 5 min. Differentiation was induced with IMDM (Invitrogen, 31980022) + 2% horse serum + 1% Penstrep. After 24 h, a thick layer of matrigel (Corning, 1:3 in differentiation medium) was added and cells were kept with 80 mg/ml of agrin.

**Table 1:**
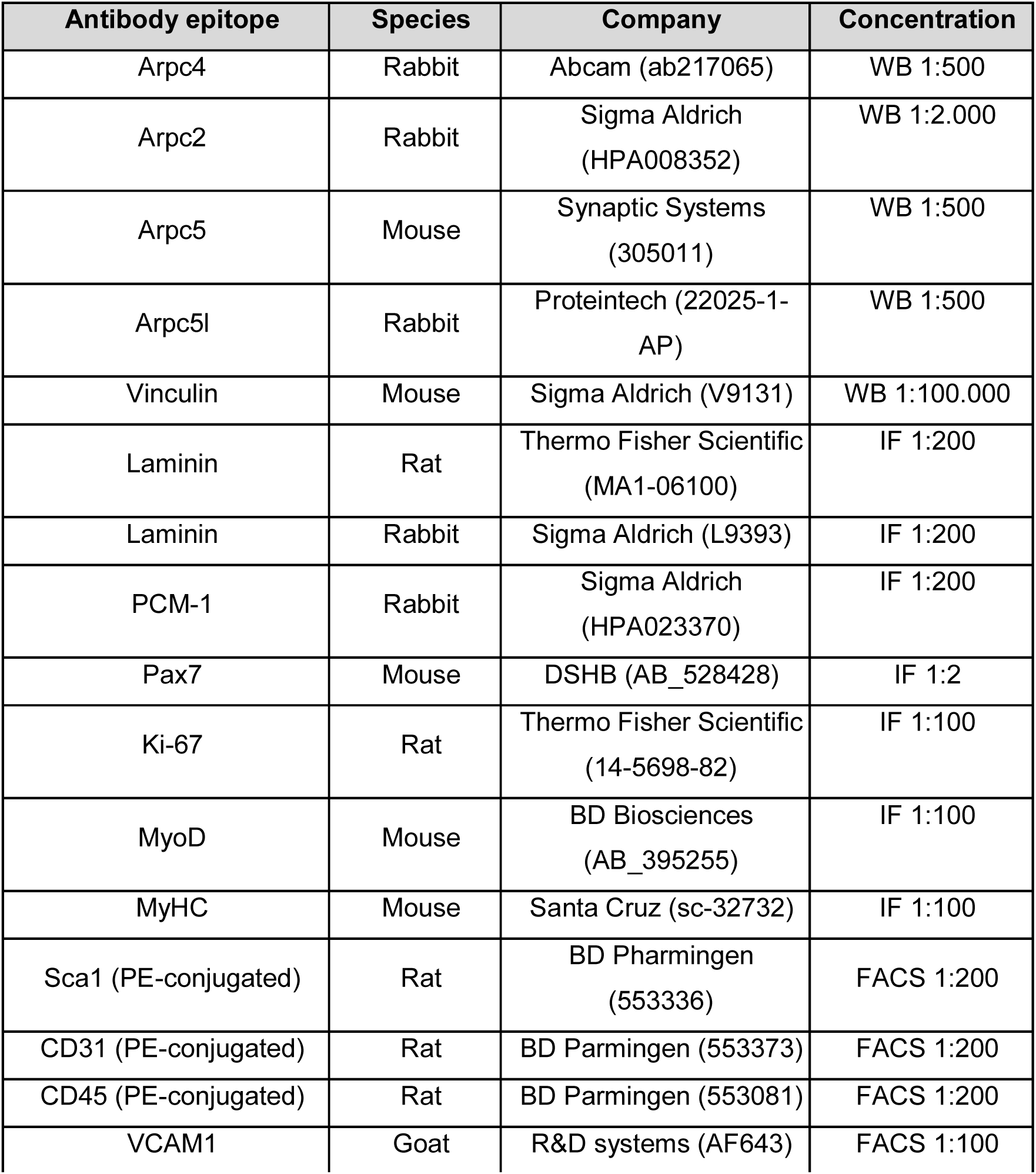
Antibodies used in this study.

| **Antibody epitope** | **Species** | **Company** | **Concentration** |
| --- | --- | --- | --- |
| Arpc4 | Rabbit | Abcam (ab217065) | WB 1:500 |
| Arpc2 | Rabbit | Sigma Aldrich<br>(HPA008352) | WB 1:2.000 |
| Arpc5 | Mouse | Synaptic Systems<br>(305011) | WB 1:500 |
| Arpc5l | Rabbit | Proteintech (22025-1-AP) | WB 1:500 |
| Vinculin | Mouse | Sigma Aldrich (V9131) | WB 1:100.000 |
| Laminin | Rat | Thermo Fisher Scientific<br>(MA1-06100) | IF 1:200 |
| Laminin | Rabbit | Sigma Aldrich (L9393) | IF 1:200 |
| PCM-1 | Rabbit | Sigma Aldrich<br>(HPA023370) | IF 1:200 |
| Pax7 | Mouse | DSHB (AB_528428) | IF 1:2 |
| Ki-67 | Rat | Thermo Fisher Scientific<br>(14-5698-82) | IF 1:100 |
| MyoD | Mouse | BD Biosciences<br>(AB_395255) | IF 1:100 |
| MyHC | Mouse | Santa Cruz (sc-32732) | IF 1:100 |
| Sca1 (PE-conjugated) | Rat | BD Pharmingen<br>(553336) | FACS 1:200 |
| CD31 (PE-conjugated) | Rat | BD Pharmingen (553373) | FACS 1:200 |
| CD45 (PE-conjugated) | Rat | BD Pharmingen (553081) | FACS 1:200 |
| VCAM1 | Goat | R&D systems (AF643) | FACS 1:100 |

### Co-culture SCs-derived myoblasts with myofibers

Myoblasts derived from isolated SCs were differentiated at 70-80% confluency with differentiation medium. After 24h, a thick layer of matrigel (Corning, 1:3 in IMDM) was added, and cells were kept with 80 mg/ml of agrin. 3 days after differentiation (D3), 20 000 cells/cm^2^ of WT proliferative SCs-derived myoblasts (unlabeled or marked with EdU) were plated on top of myotubes. After 5-6h in the growth medium to allow cell adherence, cells were kept in differentiation medium with agrin for at least 48h. Cells were either imaged in live imaging or fixed in 4% paraformaldehyde (PFA) for 10 min at room temperature (RT). Fixed co-cultures were stained for MyHC, DAPI and EdU to quantify the percentage of newly fused nuclei (Myonuclear Accretion Index, MAI).

### AAV1-CAG-CAAX-GFP production

The adenoviruses were generated by the Molecular and Transgenic Tools Platform of Fundação Champalimaud, Lisbon, following the protocols from Addgene with minor modifications ^48,49^. Briefly, production began by triple transfecting the pAAV-CAG-CAAX, pAAV-Ef1a-mCherry and pAAV-EF1a-IRES-Cre plasmid with AAV packaging plasmid expressing Rep2 and Cap1 (#112862, Addgene) and pHelper into 15 cm plates of confluent HEK-293T cells using Polyethylenimine (Polysciences Inc., catalog # 23966-100). The cells were harvested three days post-transfection. The purification step was performed by iodixanol gradient ultracentrifugation, and the viral fraction was concentrated to approximately 150 μL using an Amicon Ultra-15 centrifugal filter unit (Merck, catalog #UFC910024). For measuring virus titer, we performed qPCR (Biorad CFX96 Real-Time PCR system), using primers against Woodchuck Hepatitis Virus Posttranscriptional Regulatory Element (wpre).

### SCs transfection and infection

For Arpc4 depletion *in vitro*, cells were transfected overnight (O/N) at D0 using Lipofectamine RNAiMAX, with siRNA duplexes targeting Arpc4 (s86206 and s86207) or scramble control siRNA (4390843), according to the manufacturer’s recommendations (Thermo Fisher Scientific). For the same purpose, SCs isolated from Arpc4^fl/fl^ mice were infected at D0 with AAV9-Ef1a-mCherry (CTR) or AAV9-EF1a-IRES-Cre (Arpc4^-/-^). For the time-lapse analysis of membrane extensions, myoblasts were infected 2 days before D0 with AAV1-CAG-GFP-CAAX. The viruses were produced by Molecular and Transgenic Tools Platform (Champalimaud Foundation, Lisbon, Portugal) and used in a concentration of 3.83 x 10^6^ genome copies for 1mm^2^ well containing myoblats at approximately 70-80% confluency and incubated O/N. For the localization of the Arp2/3 complex, cells were transfected O/N at D0 using Lipofectamine 3000 (Invitrogen) according to the manufacturer’s instructions, with Arp2–mNG (kindly provided by Elizabeth Chen Lab, UT Southwestern, Dallas, Texas) ^11^. For the optogenetic activation of Cdc42, cells were co-transfected with CIBN-GFP-CAAX with either mCherry-CRY2 or mCherry-CRY2-ITSN (gifts from Mathieu Coppey, Institut Curie, Paris) O/N at D0 using Lipofectamine 3000 (Invitrogen) according to the manufacturer’s instructions ^50^.

### Western Blots

After snap-frozen in liquid nitrogen, muscle, brain or heart tissues were disrupted using Omni Soft Tissue Tip Homogenizing (Biolabproducts GmbH). Protein lysates were prepared using RIPA buffer (NaCl 150 mM, EDTA 5 mM, 1% Triton X-100, 0.5% NP-40, Tris 50 nM pH 7.5) with protease and phosphatase inhibitors (Roche). Protein quantification was performed using the BCA Protein Assay Kit (Pierce) following the manufacturer’s protocol. 20-50 ug of total protein was loaded onto 4–15% Mini-PROTEAN® TGX™ Precast Protein Gels (Bio-Rad) and transferred to Immobilon®-NC HAHY membranes (Sigma Aldrich) using a wet electroblotting system (Bio-Rad). After blocking with 4% dry milk powder, primary antibodies (Table 1) were incubated overnight at 4°C. After extensive washing with TBS-T (TBS containing 0.1% Tween-20), membranes were incubated with horseradish peroxidase (HRP)-conjugated secondary antibodies (1:5000) for 1 h at RT. For developing, membranes were incubated with Amersham ECL Prime Western Blotting Detection Reagent (Cytiva) and imaged with either Amesham 680 or Amesham 800 (GE Healthcare). Western blot densitometry was performed in unprocessed images using Image Lab 6.1.0 (Bio-Rad).

### Immunohistochemistry and image acquisition

For the measurement of myofibers’ cross-sectional area (CSA), the gastrocnemius muscle was wrapped in tragacanth gum and frozen in liquid nitrogen-cooled isopentane. For the other histological analysis, the gastrocnemius muscle was immediately fixed in 2% PFA for 2h at 4°C. After washing with PBS, the tissue was left O/N at 4°C in 15% sucrose in H_2_O. The next day, we incubated the muscle tissue in 30% sucrose in H_2_O for 3h at 4°C and then embedded it in tragacanth gum and froze it in liquid nitrogen-cooled isopentane. Transverse or longitudinal 6-10 μm cryosections were prepared from the median part of the gastrocnemius muscle with a cryostat and, after thawing to RT, Heat-Induced Epitope Retrieval (HIER) was performed in the cryosections using a PT module (A80400012, Fisher Scientific, New Hampshire, USA) at 95°C for 20 min in EnVision FLEX Target Retrieval Solution Low pH (Dako, Agilent Technologies, Denmark). After 1h of washing in PBS-T (PBS containing 0.1% Tween-20), the tissue was permeabilised, and non-specific sites were blocked with 5% BSA and 0.1% Triton X-100 in PBS for 2h at RT. Either AffiniPure™ Fab Fragment Goat Anti-Mouse (Jackson ImmunoResearch) or VisUBlock Mouse on Mouse Blocking (R&D systems) was used for the labeling of cryosections with mouse monoclonal primary antibodies. Primary antibodies (Table 1) were incubated O/N at 4°C in BSA 2%, Triton X-100 0.1% and goat serum 5% in PBS. After 1h washing in PBS-T, secondary antibodies were incubated for 2h30min at RT in BSA 2%, Triton X-100 0.1% and goat serum 5% in PBS. After another round of extensive washes with PBS-T and subsequent drying at RT, cryosections were mounted in Fluoromount G (Thermo Fisher). Tiles of the whole sections or specific positions were acquired using Zeiss Cell Observer 40x 0.75 EC Plan-NeoFluar. Tissue sectioning and the HIER were performed at the Histopathology Platform at the Gulbenkian Institute for Molecular Medicine.

### Immunocytochemistry and image acquisition

*In vitro* myofibers were fixed in 4% PFA for 10 min and, after washing in PBS, permeabilized with Triton X-100 0.2% in PBS for 20 min at RT. Non-specific sites were blocked with BSA 5% in PBS for 1 hour at RT. Primary antibodies (Table 1) were added O/N at 4°C in BSA 1% in PBS. Myofibers were washed three times and then incubated with secondary antibodies together with DAPI for 1h at RT. Myofibers were then washed 3 times with PBS before being covered with Fluoromount-G (Thermo Fisher Scientific). For the co-culture experiments, at least two 4x4 tiles were acquired per well using a Zeiss Cell Observer with a 40x 0.75 EC Plan-NeoFluar Objective.

### *In vivo* SCs fusion assay

To detect *in vivo* incorporation of myoblasts into myofibers, EdU (5-ethynyl-2′-deoxyuridine, Thermo Fisher Scientific) was administered by intraperitoneal injection (0.3 mg per 10 g of mice weight) to Arpc4^WT^ and Arpc4^-/-^ neonates 18h before muscle collection. To detect *in vitro* fusion events, we incubated cells overnight with 10 uM of EdU in growth medium. EdU detection was performed using the Click-iT™ EdU Alexa fluor 488 imaging kit as before ^16^. Myonuclear accretion to myofibers was calculated by scoring the total number of EdU+ myonuclei (EdU^+^/PCM-1^+^) per myofiber (using laminin staining).

### Image analysis

For myofiber morphometric measurements, the basal lamina was segmented using laminin staining and Cellpose (Stringer et al., 2021). The difference between myonuclei and interstitial nuclei was determined by classifying all PCM-1 positive nuclei as myonuclei, as previously established ^51,52^. Myofiber shape descriptors analyzed include circularity (4π x area/perimeter^2^), in which 1 indicates a myofiber whose shape is a perfect circle and roundness (4 x area/π x major_axis^2^), which considers the relationship between myofiber area and its longest axis. The fusion index was calculated by dividing the number of nuclei inside myotubes with more than 2 nuclei by the total number of nuclei in the field of view.

### Time-lapse imaging and analysis

To analyze myofiber membrane dynamics in contact with myoblasts, time-lapse imaging was performed in Visitech VT-iSIM microscope at 37 °C in 5% CO_2_. Myofibers were infected with an AAV1 carrying CAAX-GFP and differentiated for 3 days. At D3, cells were co-cultured with tdTomato-myoblasts in growth medium for at least 4 hours, and then changed to differentiation medium. Imaging started more than 15h after changing to the differentiation medium, to allow the cells to stabilize. Then, we acquired every 5 min for 24-48 h, using a Nikon Plan Apo 40x/1.0 Oil Ph3 DM objective. Membrane extensions were counted per 50 µm of the myofiber cell cortex, in regions where the cell was in contact with a myoblast for more than 1 h.

To follow Arp2/3 localization during fusion, time-lapse imaging was performed using Zeiss LSM980 with Airyscan 2 microscope at 37 °C in 5% CO_2_. Myofibers were transfected with Arp2-mNeonGreen (Arp2-mNG) and differentiated for 3 days. At D3, cells were co-cultured with unlabeled SCs in growth medium for at least 4 hours, and then changed to differentiation medium. Imaging started more than 15h after changing to the differentiation medium, to allow the cells to stabilise. Then, we acquired every 5 min for 30h, using a 63x Plan-Apochromat DIC Oil 1.40 Objective. To study the Arp2/3 accumulation in myofibers interacting with myoblasts, images were acquired in the Visitech VT-iSIM, using a CFI SR HP Apo TIRF 100XAC Oil 1.49 Objective. We analyzed the presence of Arp2-mNG, the contact with a neighbouring cell and the presence of membrane extensions at every 20 µm of the myofiber cell cortex.

For the optogenetic activation of Cdc42, time-lapse imaging was performed using Zeiss LSM980 with Airyscan 2 microscope at 37 °C in 5% CO_2_. Myofibers were co-transfected with CIBN-GFP-CAAX and either mCherry-CRY2 or mCherry-CRY2-ITSN, and differentiated for 3 days. At D3-D4, cells were co-cultured with unlabeled SCs in growth medium for at least 4 hours, and then changed to differentiation medium. We used 488 nm light at 20% laser power to induce optogenetic activation of the CIBN/CRY2 pair every 6 min for 12h, using a 63x Plan-Apochromat DIC Oil 1.40 Objective.

All microscopy experiments were performed at the Bioimaging Platform of the Gulbenkian Institute for Molecular Medicine

### Data representation and statistics

All graphs were plotted using GraphPad Prism (version 8.4.3 of GraphPad Software Inc.), and figures were formatted with InkScape (version 1.2.2). Statistical analysis was performed with GraphPad Prism. Parametric or nonparametric tests were selected based on whether the samples were normally distributed. Statistical significance was determined by unpaired 2-tailed Student’s t test, unpaired 2-tailed Mann-Whitney test, one-way ANOVA with Dunnett’s post-hoc test or two-way ANOVA with Sidak’s post-hoc test, as indicated in the figure legends. Differences between groups were considered statistically significant when p≤0.05. Data distributions are expressed as mean ± SEM.

## SUPPLEMENTARY FIGURES

**Supplementary Figure 1.**
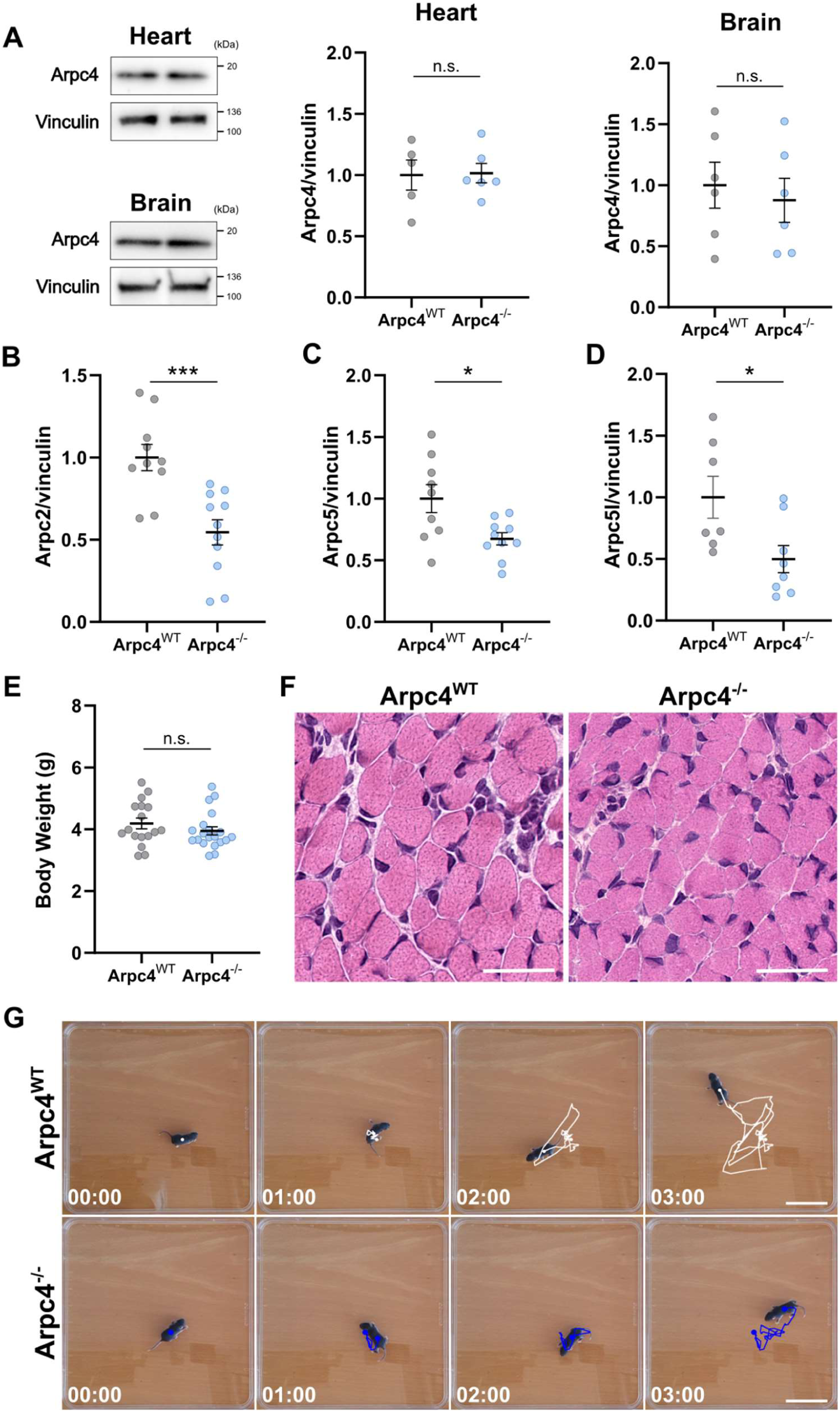
Characterization of Arpc4^-/-^ mice. **A)** (Left) Western blots (WB) for Arpc4 and Vinculin in the heart and brain of P7 Arpc4^WT^ and Arpc4^-/-^ mice. (Right) Quantification of Arpc4 protein levels by WB normalized against Vinculin, in heart and brain, as fold vs the average of littermate Arpc4^WT^. **B-D)** Quantification of WB from Fig. 1B, showing the protein levels of Arpc2 (B), Arpc5 (C), and Arpc5l (D) normalized against Vinculin. The results are shown as fold vs. the average of littermate Arpc4^WT^. **E)** Body weight of P7 Arpc4^WT^ and Arpc4^-/-^ mice. Data are represented as mean ± s.e.m. (n≥5 mice). *p≤0.05, **p≤0.01, ***p≤0.001 using unpaired 2-tailed Student’s t test. **F)** Representative images of cross-sectioned gastrocnemius muscle from P7 Arpc4^WT^ and Arpc4^-/-^ mice, stained for Hematoxylin and Eosin (H&E). **G)** Representative frames of a time lapse movie with the trajectories of a P7 Arpc4^WT^ (white) and Arpc4^-/-^ (blue) mice during 3 min on an open field, showing min 0, 1, 2 and 3 separately. Scale bar: 5 cm.

**Supplementary Figure 2.**
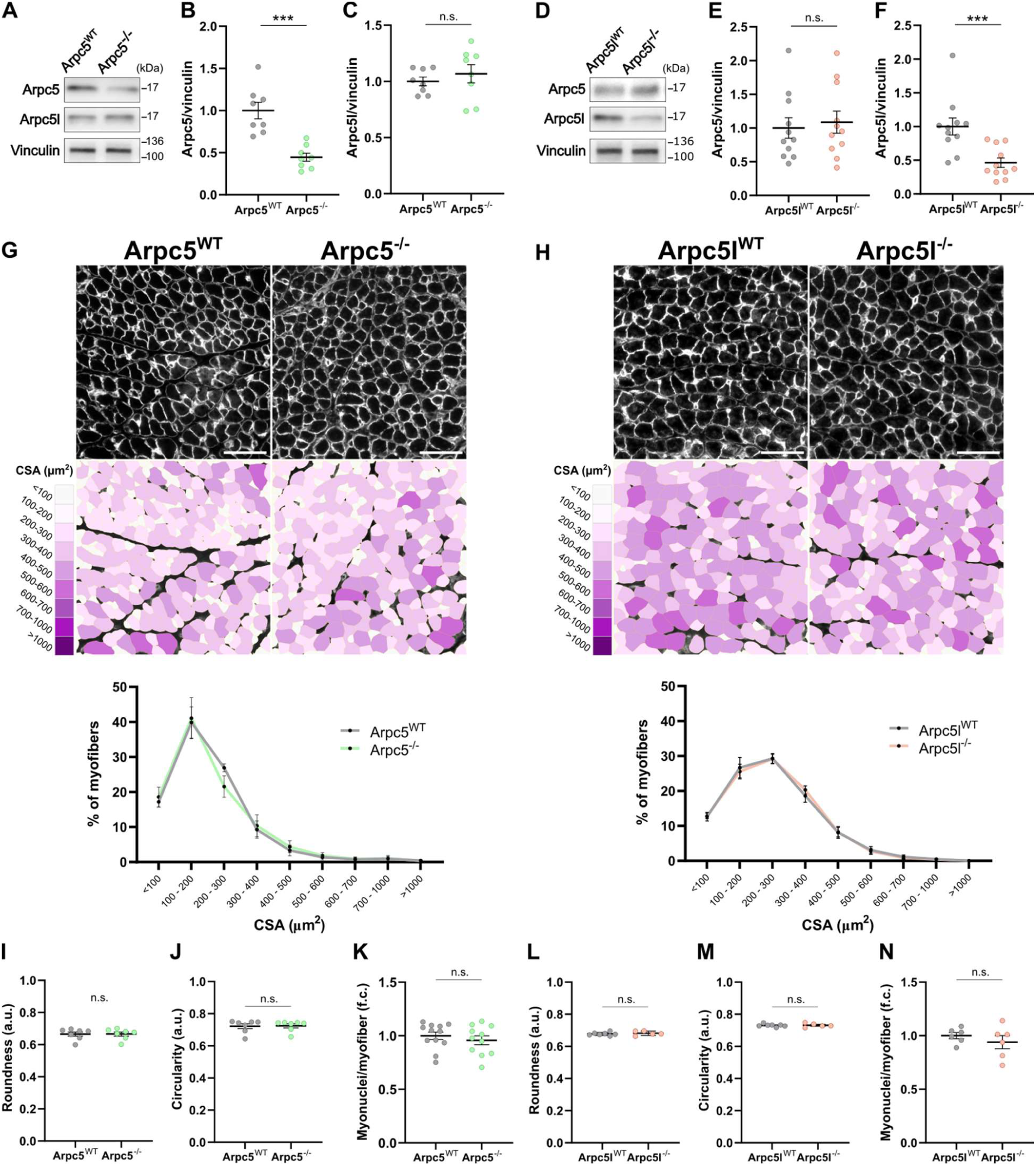
The role of the Arp2/3 complex in myofiber size and nuclear number is not Arpc5 isoform-dependent. **A)** Representative WB for Arpc5, Arpc5l and Vinculin from skeletal muscle lysate of P7 Arpc5^WT^ and Arpc5^-/-^ mice. **B-C)** Quantification of (B) Arpc5 and (C) Arpc5l protein levels by WB, shown in (A), normalized against Vinculin. The results are shown as fold vs. the average of littermate Arpc5^WT^. **D)** Representative WB for Arpc5, Arpc5l and Vinculin from skeletal muscle lysate of P7 Arpc5l^WT^ and Arpc5l^-/-^ mice. **E,F)** Quantification of (E) Arpc5 and (F) Arpc5l protein levels by WB, shown in (D), normalized against Vinculin. The results are shown as fold vs. the average of littermate Arpc5l^WT^. **G,H)** Representative images of the P7 (G) Arpc5^WT^/Arpc5^-/-^ and (H) Arpc5l^WT^/Arpc5l^-/-^ mice gastrocnemius muscle. (Top) cross-sections stained for the basal lamina (laminin, white). (Middle) top images, colour-coded by CSA, as indicated in the legend. Scale bar: 50 µm. (Bottom) Frequency distribution of myofiber CSA. Data are represented as mean ± s.e.m. (n≥5 mice). **I-K)** Quantifications of the myofiber (K) roundness, (L) circularity and (K) myonuclei number in P7 Arpc5^WT^ and Arpc5^-/-^ mice myofibers. **L-M)** Quantifications of the myofiber (L) roundness, (M) circularity and (N) myonuclei number in P7 Arpc5l^WT^ and Arpc5l^-/-^ mice myofibers. The number of myonuclei per myofiber is expressed as fold vs. the average of littermate controls. Data are represented as mean ± s.e.m. (n≥5 mice). *p≤0.05, **p≤0.01, ***p≤0.001 using unpaired 2-tailed Student’s t test or unpaired 2-tailed Mann-Whitney test, depending on normal distribution, in all except G and H, where two-way ANOVA with Sidak’s post-hoc test was used.

**Supplementary Figure 3.**
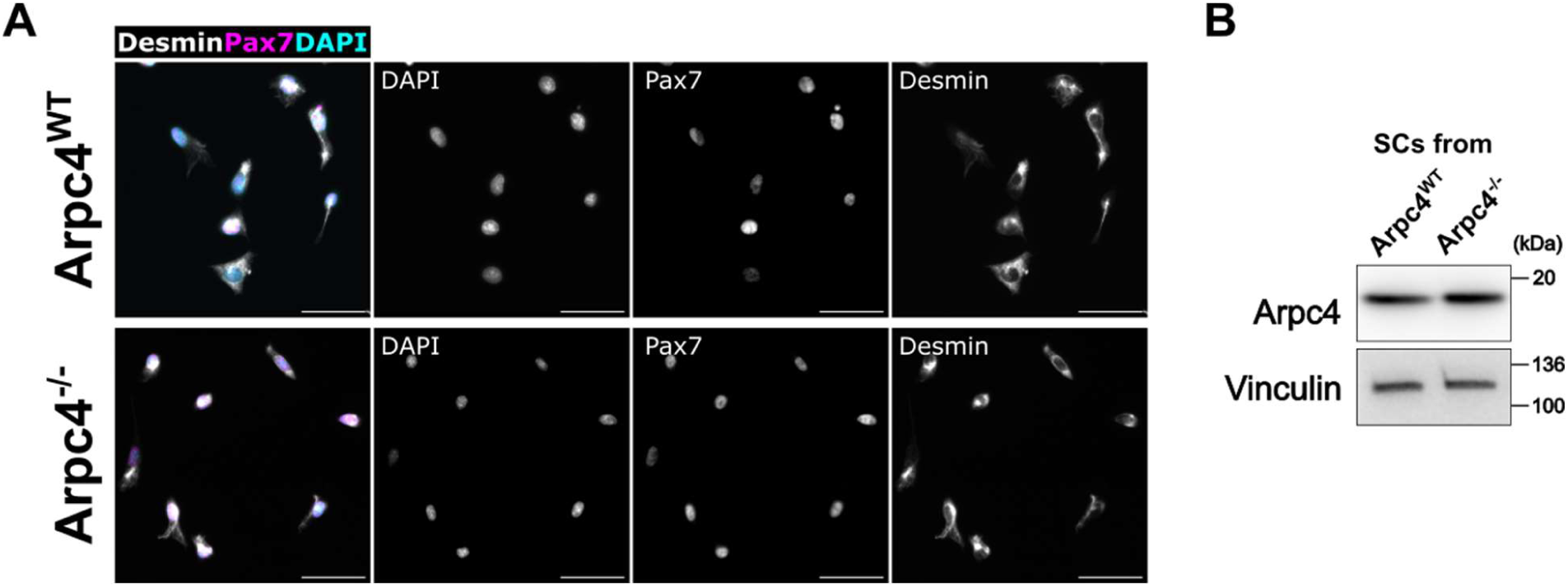
Characterization of SCs isolated via FACS-sorting. **A)** Representative image of the population obtained from SCs isolated from the skeletal muscle of P7 Arpc4^WT^ and Arpc4^-/-^ mice, stained for Desmin (white), Pax7 (magenta) and DAPI (cyan). Scale bar: 50 µm. **B** Western blot for Arpc4 and Vinculin in SCs sorted from P7 Arpc4^WT^ and Arpc4^-/-^ mice.

**Supplementary Figure 4.**
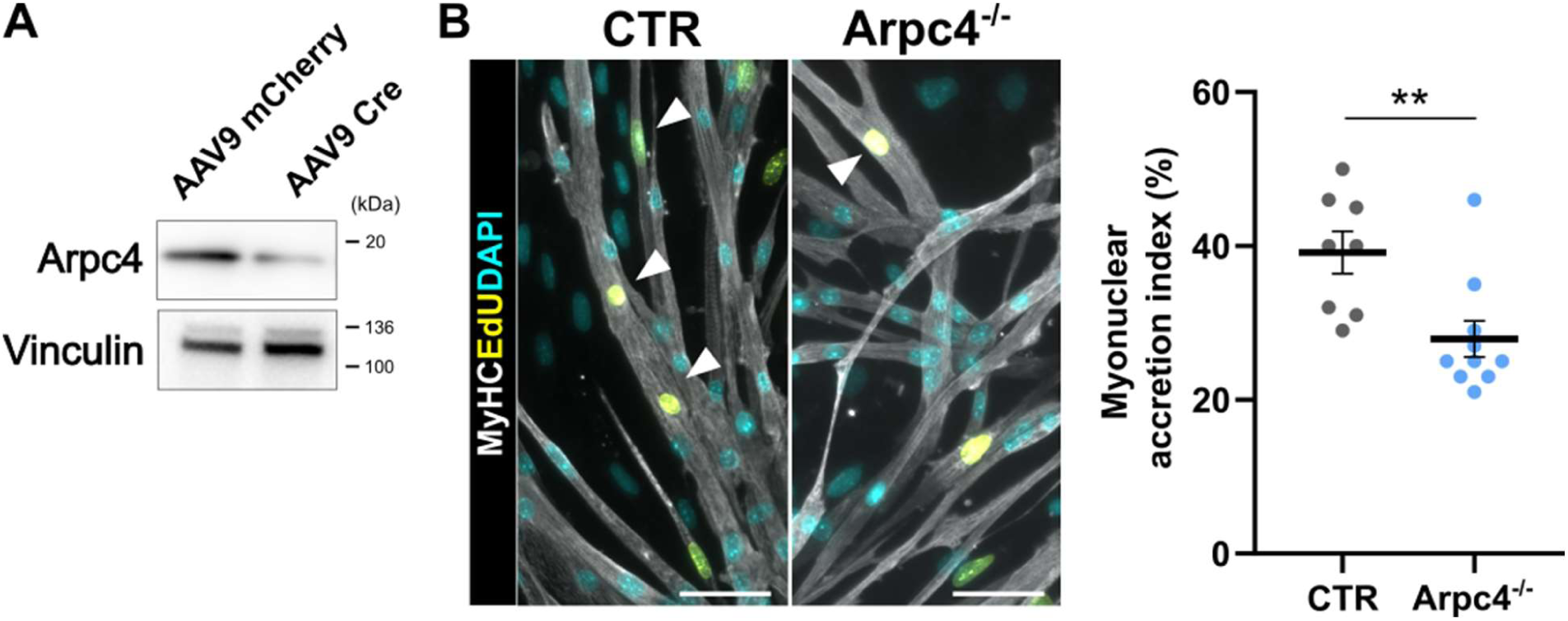
Myoblasts failed to fuse to Arpc4-depleted myofibers *in vitro*. **A)** Western blots for Arpc4 and Vinculin in D3 myotubes generated from Arpc4^fl/fl^ SCs infected with an AAV9 carrying Cre recombinase. **B)** (Left) Representative images of the co-culture between D3 myotubes and EdU-labeled WT myoblasts, stained for MyHC (white), EdU (magenta) and DAPI (cyan). White arrows identify EdU+ myonuclei in pre-existing myofibers. Scale bar: 50 µm. (Right) Quantification of the myonuclear accretion index. Data are represented as mean ± s.e.m. (n≥7 independent experiments). *p≤0.05, **p≤0.01, ***p≤0.001 using (F) one-way ANOVA with Dunnett’s post-hoc test or (H) unpaired 2-tailed Mann-Whitney test.

## MOVIES

**Sup. Movie 1:** Time-lapse movie of a myofiber creating long-lived membrane protrusions upon interaction with myoblasts (magenta). Myofiber membrane is labeled with a CAAX-GFP (white). Time in hh:min; Scale bar: 20 µm.

**Sup. Movie 2:** Time-lapse movie showing Arp2-mNG (white) in a myofiber during interaction with unlabeled myoblasts. In the BF, it is possible to follow the position of the unlabeled myoblast relative to the myofiber. Fusion is observed at 11:25 (hh:min). Time in hh:min; Scale bar: 20 µm.

**Sup. Movie 3:** Time-lapse movie of a mCherry^Opto^ myofiber (white) interacting with an unlabeled myoblast. Myofiber membrane is labeled with a CAAX-GFP (white). In the BF, it is possible to follow the position of the unlabeled myoblast relative to the myofiber. Time in hh:min; Scale bar: 20 µm.

**Sup. Movie 4:** Time-lapse movie of a Cdc42^Opto^ myofiber (white) interacting with an unlabeled myoblast. Myofiber membrane is labeled with a CAAX-GFP (white). In the BF, it is possible to follow the position of the unlabeled myoblast relative to the myofiber. Fusion is observed at 10:06 (hh:min). Time in hh:min; Scale bar: 20 µm.

**Sup. Movie 5:** Time-lapse movie of a Arpc4^WT^ myofiber interacting with a myoblast (magenta). Myofiber membrane is labeled with a CAAX-GFP (white). Time in hh:min; Scale bar: 20 µm.

**Sup. Movie 6:** Time-lapse movie of a Arpc4^-/-^ myofiber interacting with a myoblast (magenta). Myofiber membrane is labeled with a CAAX-GFP (white). Time in hh:min; Scale bar: 20 µm.

